# Microglia produce the amyloidogenic ABri peptide in familial British dementia

**DOI:** 10.1101/2023.06.27.546552

**Authors:** Charles Arber, Jackie M. Casey, Samuel Crawford, Naiomi Rambarack, Umran Yaman, Sarah Wiethoff, Emma Augustin, Thomas M. Piers, Agueda Rostagno, Jorge Ghiso, Patrick A. Lewis, Tamas Revesz, John Hardy, Jennifer M. Pocock, Henry Houlden, Jonathan M. Schott, Dervis A. Salih, Tammaryn Lashley, Selina Wray

## Abstract

Mutations in *ITM2B* cause familial British, Danish, Chinese and Korean dementias. In familial British dementia (FBD) a mutation in the stop codon of the *ITM2B* gene (also known as *BRI2*) causes a C-terminal cleavage fragment of the ITM2B/BRI2 protein to be extended by 11 amino acids. This fragment, termed amyloid-Bri (ABri), is highly insoluble and forms extracellular plaques in the brain. ABri plaques are accompanied by tau pathology, neuronal cell death and progressive dementia, with striking parallels to the aetiology and pathogenesis of Alzheimer’s disease. The molecular mechanisms underpinning FBD are ill-defined. Using patient-derived induced pluripotent stem cells, we show that expression of *ITM2B/BRI2* is 34-fold higher in microglia than neurons, and 15-fold higher in microglia compared with astrocytes. This cell-specific enrichment is supported by expression data from both mouse and human brain tissue. ITM2B/BRI2 protein levels are higher in iPSC-microglia compared with neurons and astrocytes. Consequently, the ABri peptide was detected in patient iPSC-derived microglial lysates and conditioned media but was undetectable in patient-derived neurons and control microglia. Pathological examination of post-mortem tissue support ABri expression in microglia that are in proximity to pre-amyloid deposits. Finally, gene co-expression analysis supports a role for ITM2B/BRI2 in disease-associated microglial responses. These data demonstrate that microglia are the major contributors to the production of amyloid forming peptides in FBD, potentially acting as instigators of neurodegeneration. Additionally, these data also suggest ITM2B/BRI2 may be part of a microglial response to disease, motivating further investigations of its role in microglial activation. This has implications for our understanding of the role of microglia and the innate immune response in the pathogenesis of FBD and other neurodegenerative dementias including Alzheimer’s disease.

## Introduction

Mutations in *ITM2B* (also known as *BRI2*) cause familial British (Vidal et al., 1999), Danish (Vidal et al., 2000), Chinese (Liu et al., 2021) and Korean (Rhyu et al., 2023) dementias (FBD, FDD, FCD and FKD respectively) and have been associated with autosomal dominant retinal dystrophy (Audo et al., 2014). ITM2B/BRI2 is a type II transmembrane protein that is cleaved by FURIN convertase to release an extracellular 23 amino acid C terminal fragment (Kim et al., 1999; Seong-Hun et al., 2000). Dominantly inherited mutations linked to dementia increase the length of this C terminal fragment to 34 amino acids, either by disrupting the stop codon (in FBD, FCD and FKD), or due to a 10 nucleotide duplication upstream of the stop codon (in FDD). These extended peptides are amyloidogenic and termed Amyloid Bri (ABri) in FBD and Amyloid Dan (ADan) in FDD; no histopathological data are available for FCD or FKD to date, but the mutations are predicted to produce a peptide that differs from ABri by a single amino acid. These amyloidogenic peptides aggregate to form extracellular amyloid deposits (Ghiso et al., 2000), and are thus theorised to cause neurodegeneration in a manner analogous to Amyloid-beta (Aβ) in Alzheimer’s disease (AD) (Del Campo and Teunissen, 2014; Mead et al., 2000).

Common clinical features of FBD and FDD include progressive dementia, and cerebellar ataxia (Worster-Drought et al., 1933). Additional specific features include spastic paraparesis in FBD, as well as cataracts and hearing defects in FDD (Liu et al., 2021; Plant et al., 1990; Strömgren et al., 1970). Pathologically, FBD and FDD are characterised by extensive amyloid angiopathy, amyloid plaques and pre-amyloid deposits in multiple brain regions. ABri and ADan also accumulate in organs throughout the body (Ghiso et al., 2001). Similar to AD, and supporting shared aetiology, neurofibrillary tangle tau pathology is common in both FBD and FDD (Holton et al., 2002, 2001).

The normal function of ITM2B/BRI2 is not well understood. The protein is localised to the plasma membrane and potentially the mitochondrial inner membrane (Wohlschlegel et al., 2021) and has been reported to interact with APP and Aβ - modulating Aβ deposition (Fotinopoulou et al., 2005; Kim et al., 2008; Tamayev et al., 2010). The BRICHOS domain, which is distinct from the cleaved C-terminus fragment, is able to reduce the fibrillation of amyloids (Peng et al., 2010; Willander et al., 2012). Conversely, APP was shown to be a molecular effector of ADan-associated synaptic and memory deficits, as APP haploinsufficiency prevents synaptic and memory deficits in a mouse model of FDD (Tamayev et al., 2010). Aβ co-accumulates with ADan pathology but this has not been observed with ABri (Holton et al., 2002; Michno et al., 2022; Tomidokoro et al., 2005). Mechanistically, there remains uncertainty whether mutations cause a loss of normal function of the ITM2B/BRI2 protein, as knock-in mouse models have shown reduced expression of ITM2B/BRI2 protein (Tamayev et al., 2010; Yin et al., 2021).

Genome wide association studies have revealed the importance of microglia in the determination of risk for several neurodegenerative disorders, particularly Alzheimer’s disease (Podleśny-Drabiniok et al., 2020; Villegas-Llerena et al., 2016). Further, heterozygous mutations in *TREM2*, a microglial gene, increase risk for AD three-fold (Guerreiro et al., 2013; Jonsson et al., 2013). The molecular mechanism by which microglia contribute to dementia onset and progression is an area of intense investigation. Recent work suggests that microglia regulate the transition of amyloid pathology to tau pathology (Lee et al., 2021; Leyns et al., 2019) via genes expressed by the amyloid-responsive microglial state (ARM) or the disease-associated microglial state (DAM) (Keren-Shaul et al., 2017; Matarin et al., 2015; Nguyen et al., 2020; Salih et al., 2019); cell states that are enriched in dementia and driven by genes including *APOE* and *TREM2*.

In this study, we sought to gain insights into the cellular consequences of FBD-associated *ITM2B/BRI2* mutations by developing patient-derived iPSC models of FBD. Based on previous pathological findings that ITM2B/BRI2 can be expressed by neurons and glia (Lashley et al., 2008), we sought to determine the effect of FBD mutations on ITM2B/BRI2 in different cell types. Due to the parallels between FBD, FDD and AD, and the crucial role of microglia in AD progression (Hardy and Salih, 2021; Hodges et al., 2021; Podleśny-Drabiniok et al., 2020), we then investigated a possible role of microglia in FBD and FDD pathobiology.

## Results

To generate a human, physiological model of FBD, we reprogrammed fibroblasts from two individuals with the FBD mutation, c.799 T>A (for donor details, see Table 1). iPSCs showed characteristic morphology and expression of the pluripotency markers SSEA1, OCT4, SOX2, NANOG and DNMT3B via immunocytochemistry and qPCR (Fig 1A-B). Reprogrammed cells also showed appropriate downregulation of the fibroblast specific markers *S100A4* and *VIM* (Fig 1C). FBD iPSCs were further characterised to confirm the presence of the FBD mutation TGA>AGA (Fig 1D), as well as a stable karyotype (Fig 1E). Analysis of a panel of 770 genes associated with pluripotency and early differentiation demonstrated that the two FBD iPSC lines showed a global expression profile comparable to a panel of 3 control stem cell lines (Fig 1F).

**Figure 1.**
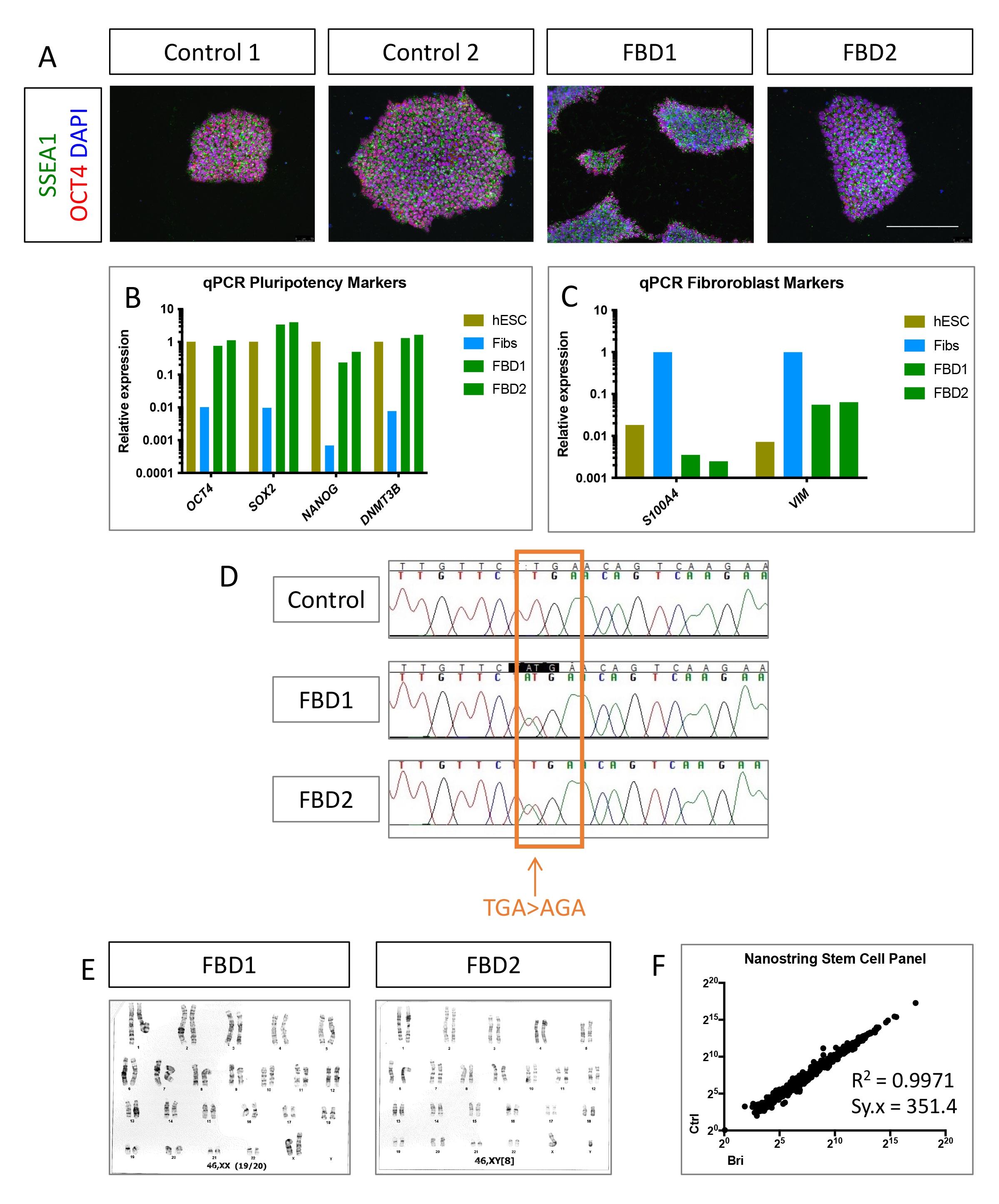
Generation and characterisation of FBD patient-derived iPSCs. A) Immunocytochemistry showing pluripotency markers (SSEA4 and OCT4) in patient-derived iPSCs (FBD1 and FBD2) and control iPSCs (Control 1-2). Scale bar represents 200μm. B-C) qPCR analysis of FBD iPSCs compared to cDNA from fibroblasts (Fibs) and control human embryonic stem cells (hESCs). Stem cell markers (*OCT4*, *NANOG*, *SOX2* and *DNMT3B*) are expressed to a similar degree in reprogrammed iPSCs and control hESCs. Fibroblast markers (*S100A4* and *VIM*) are downregulated compared with fibroblasts. D) Sanger sequencing to confirm the presence of the FBD-associated mutation in FBD iPSCs. E) Karyotype analysis demonstrated appropriate G-banding and a stable karyotype. F) Nanostring Stem Cell Characterisation panel analysis supports stable iPSC identity of FBD1 and FBD2 iPSCs when compared with three control iPSC/hESCs (Ctrl1, Ctrl2 and Shef6).

**Table 1.** Details of stem cell lines.

|  | **Gender** | **Age at biopsy** | **FBD status** | **Source** |
| --- | --- | --- | --- | --- |
| **FBD fibroblast donor 1 (FBD1)** | **Female** | **50** | **FBD - Affected** | **UCL DRC** |
| **FBD fibroblast donor 2 (FBD2)** | **Male** | **58** | **FBD - Affected** | **NHNN** |
| **Ctrl1 (RBi001-a)** | **Male** | **45-49** | **Unaffected** | **Sigma Aldrich/EBiSC** |
| **Ctrl2 (SIGi1001-a-1)** | **Female** | **20-24** | **Unaffected** | **Sigma Aldrich/EBiSC** |
| **hESC (Shef6)** | **Female** | **0** | **Unaffected** | **UK Stem Cell Bank** |

Abbreviations: FBD – familial British dementia; DRC – Dementia Research Centre; NHNN – National Hospital for Neurology and Neurosurgery.

We investigated the cell type specific expression of *ITM2B/BRI2* in control iPSC-derived neurons, astrocytes and microglia (Fig 2A-B). Successful differentiation of iPSCs to cortical neurons was confirmed by morphology and expression of neuronal-specific TUJ1 and deep layer cortical neuronal marker TBR1. IBA1 expression confirmed the successful differentiation of iPSCs to microglia-like cells. Astrocyte differentiation was confirmed via enrichment of SOX9 by Western blotting (Fig 2C). qPCR analysis demonstrated that expression of *ITM2B/BRI2* was on average 34-fold higher in microglia when compared with neurons (Fig 2B) and 14-fold higher in microglia compared with astrocytes. This cell-type specific enrichment is supported when mining expression data from mouse and human brain tissue in 7 freely available datasets (Friedman et al., 2018; Guttenplan et al., 2020; Hochgerner et al., 2018; Lake et al., 2018; Mancarci et al., 2017; Saunders et al., 2018; Zhang et al., 2016, 2014a) (Fig S1-S2). *FURIN*, encoding the enzyme responsible for cleavage of ITM2B/BRI2 in normal physiology and release of ABri and ADan in disease, showed enrichment in microglia and astrocytes relative to neurons (Fig 2B). Western blotting of neuronal, astrocytic and microglial lysates reinforced the finding that ITM2B/BRI2 is highly expressed by microglia (Fig 2C-D). The specificity of bands at around 30kDa and 12kDa, corresponding to full-length and cleaved ITM2B/BRI2 protein respectively, were confirmed by siRNA knockdown of *ITM2B/BRI2* in iPSC-derived microglia (Fig 2C and Fig S3). This finding is reinforced by a published proteomics study that shows relative depletion of ITM2B/BRI2 protein in acutely isolated mouse neurons compared with glia (Sharma et al., 2015) (Fig S4). We also detected a band at around 25kDa in human brain samples (Fig 2E); however, we cannot determine if this band is specific or is a cleavage fragment of ITM2B/BRI2.

**Figure 2.**
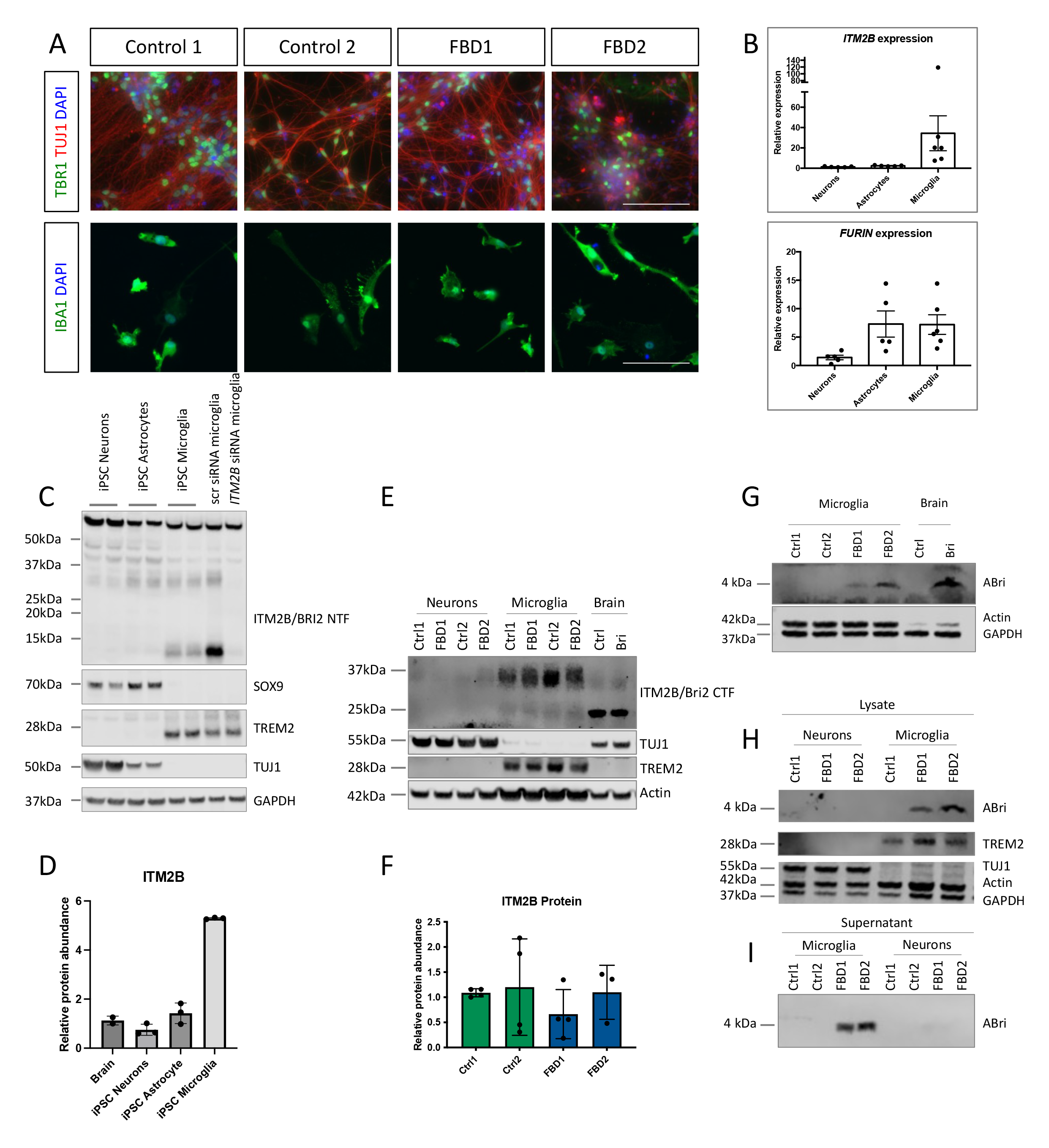
ABri is produced by iPSC-derived microglia. A) Immunocytochemistry of iPSC-derived neurons (upper panels) and microglia (lower panels). TUJ1 is a pan-neuronal marker and TBR1 labels deep layer cortical neurons, scale bar 200μm. IBA1 labels microglia-like cells, scale bar 50μm. B) qPCR analysis of *ITM2B/BRI2* and *FURIN* expression in control iPSC-derived cortical neurons, astrocytes and microglia. Neuronal cDNA represents 5 independent inductions with 2 independent control iPSC lines, astrocytic cDNA was generated from 5 independent inductions of 2 independent control iPSC lines and microglial cDNA was generated from 6 harvests from 4 inductions and represents 2 independent control iPSC lines. C) Western blotting of iPSC-derived neurons, iPSC-derived astrocytes and iPSC-derived microglia. *ITM2B/BRI2* knockdown via siRNA depicts antibody specificity for bands at around 30kDa and 12kDa. TUJ1, SOX9 and TREM2 are markers for neurons, astrocytes and microglia respectively. Samples represent two independent control lines for each cell type D) Quantification of specific bands (30kDa and 12kDa) from 3 independent neuronal, astrocyte and microglia inductions of at least two control lines in each cell type. E) Western blotting of iPSC neurons, iPSC microglia and post-mortem brain tissue for ITM2B/BRI2 as well as neuronal TUJ1, microglial TREM2 and loading control (Actin). F) Quantification of ITM2B/BRI2 in control and patient-derived microglia from 4 harvests from three independent batches of microglia. G-H) Western blotting for ABri in iPSC-derived microglia lysates and brain tissue showed a band of 4kDa. I) Western blotting for ABri in iPSC-derived microglial conditioned media.

FBD and FDD mutations have been shown to reduce the levels of ITM2B/BRI2 protein in mouse knock-in models (Tamayev et al., 2010; Yin et al., 2021). To determine if the FBD mutation affects protein levels in iPSC-derived microglia, we performed Western blotting and observed comparable protein levels in patient-derived microglia and control microglia (Fig 2E-F), albeit with a degree of variability between microglial batches.

Western blotting using the ABri-specific antibody (Ab338) was able to detect the presence of the 4kDa ABri peptide in patient-derived microglial lysates (Fig 2G-H) as well as in patient-derived microglial-conditioned media (Fig 2I). We were not able to detect the ABri peptide in FBD neuronal lysates or control microglia (Fig 2G-I).

Based on the finding that *ITM2B/BRI2* expression is enriched in microglia, we sought to investigate the pathological contribution of microglia in mutant *ITM2B/BRI2*-associated post-mortem hippocampal tissue (for donor details, see Table 2). Due to the availability of the rare FBD and FDD post-mortem tissue, we investigated both diseases which have ITM2B/BRI2- associated pathology. Analyses employed antibodies against ABri (Ab338) and ADan (Ab5282) in conjunction with microglia markers and Thioflavin staining for amyloid structures. In FBD, ABri was found in the form of amyloid and pre-amyloid plaques as previously documented (Holton et al 2001) (Fig 3A-C). Microglia were found within the amyloid plaques, but also in areas containing pre-amyloid ABri deposits (Fig 3B). The pre-amyloid diffuse ABri deposits also contained microglia shaped cells stained positive by the ABri antibody (Fig 3C, arrows). In regions of pre-amyloid deposition, fluorescent ABri immunohistochemistry (IHC) was undertaken with Thioflavin staining to confirm that the cells contained ABri in an amyloid conformation (Fig 3 row D and E, arrows). Double IHC confirmed that the amyloid was present within microglia using CD68 (Fig 3 row F) and CR3/43 (Fig 3 row G) microglial markers. Due to the availability of FDD post-mortem tissue, we further explored ITM2B/BRI2-associated amyloid in a case of FDD. ADan was found predominantly as pre-amyloid deposits (Fig 4A-B) and in the form of cerebral amyloid angiopathy. ADan immunohistochemistry clearly outlines microglia shaped cells (Fig 4C, arrows). When visualised with Thioflavin and ADan, microglia morphologies were clearly visible surrounded by the pre-amyloid deposits (Fig 4 rows D and E, arrows). Double IHC for microglial markers and Thioflavin, clearly showed the microglia containing Thioflavin positive amyloid (Fig 4 rows F and G, arrows). Taken together, these findings suggest that ABri and ADan are found in an amyloidogenic conformation within the microglia when in close proximity to pre-amyloid deposits.

**Figure 3.**
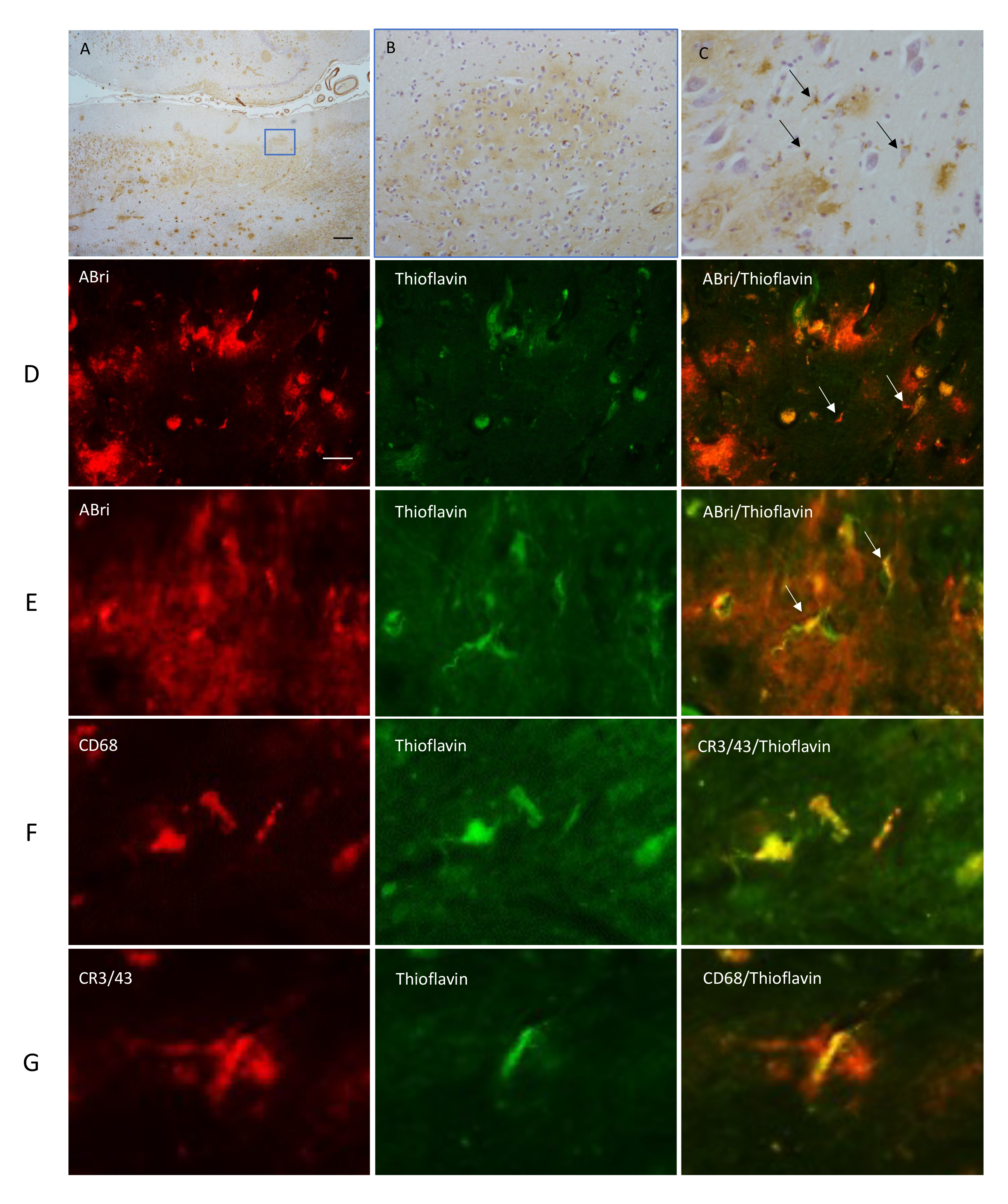
Immunohistochemical staining in FBD for ABri, Thioflavin and microglial markers. ABri pathology is observed in the hippocampus (A) in the form of extracellular amyloid and preamyloid deposits. ABri is also found within blood parenchymal and leptomeningeal blood vessels as cerebral amyloid angiopathy. ABri pre-amyloid plaques are shown at higher magnification in (B). The preamyloid deposits contained cells resembling microglia morphology (C, arrows). The bar represents 500µm in A and 50µm in B and 25µm in C. ABri immunohistochemistry (red, row D and E) combined with Thioflavin staining highlights the presence of ABri in cells resembling microglia. Microglial antibodies were used to determine the Thioflavin positive structures identified in the cells (rows F and G). The bar represents 100µm in row D and 20µm in row E and 10µm rows in F and G.

**Figure 4.**
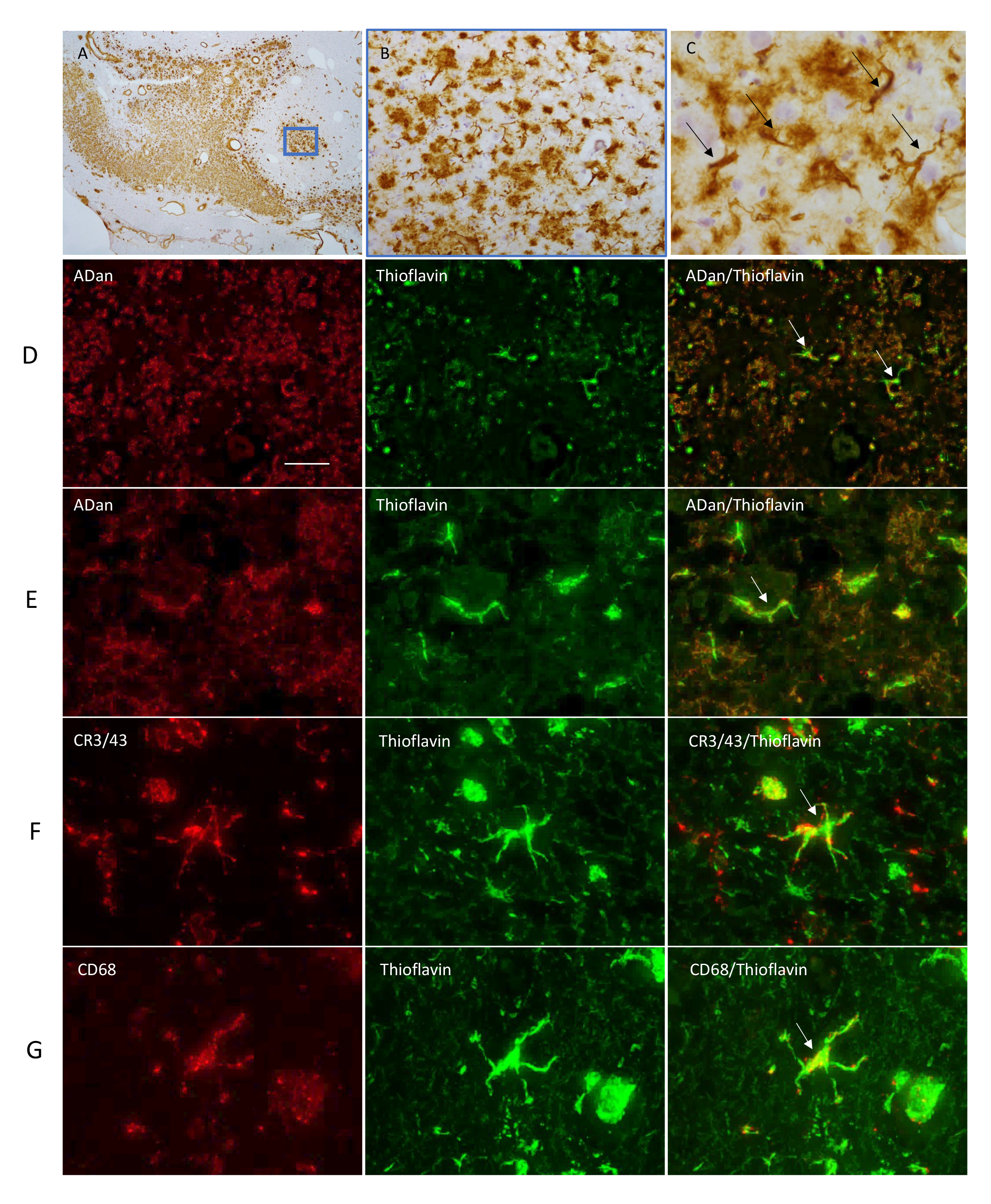
Immunohistochemical staining in FDD for ADan, Thioflavin and microglial markers. ADan pathology was observed in the hippocampus (A) in the form of extracellular pre-amyloid deposits and cerebral amyloid angiopathy. At higher magnification we observed the preamyloid deposits (B). Structures resembling microglia are also found to be highlighted with the ADan immunohistochemical preparations in the pre-amyloid deposits (C, arrows). The bar represents 500µm in A and 50µm in B and 25µm in C. ADan immunohistochemistry (red, row D and E) combined with Thioflavin staining (green) highlights the presence of ADan in cells resembling microglia. Microglial antibodies were used to determine the Thioflavin positive structures identified in the cells (rows F and G, arrows). The bar represents 100µm in row D and 20µm in row E and 10µm rows in F and G.

**Table 2.**
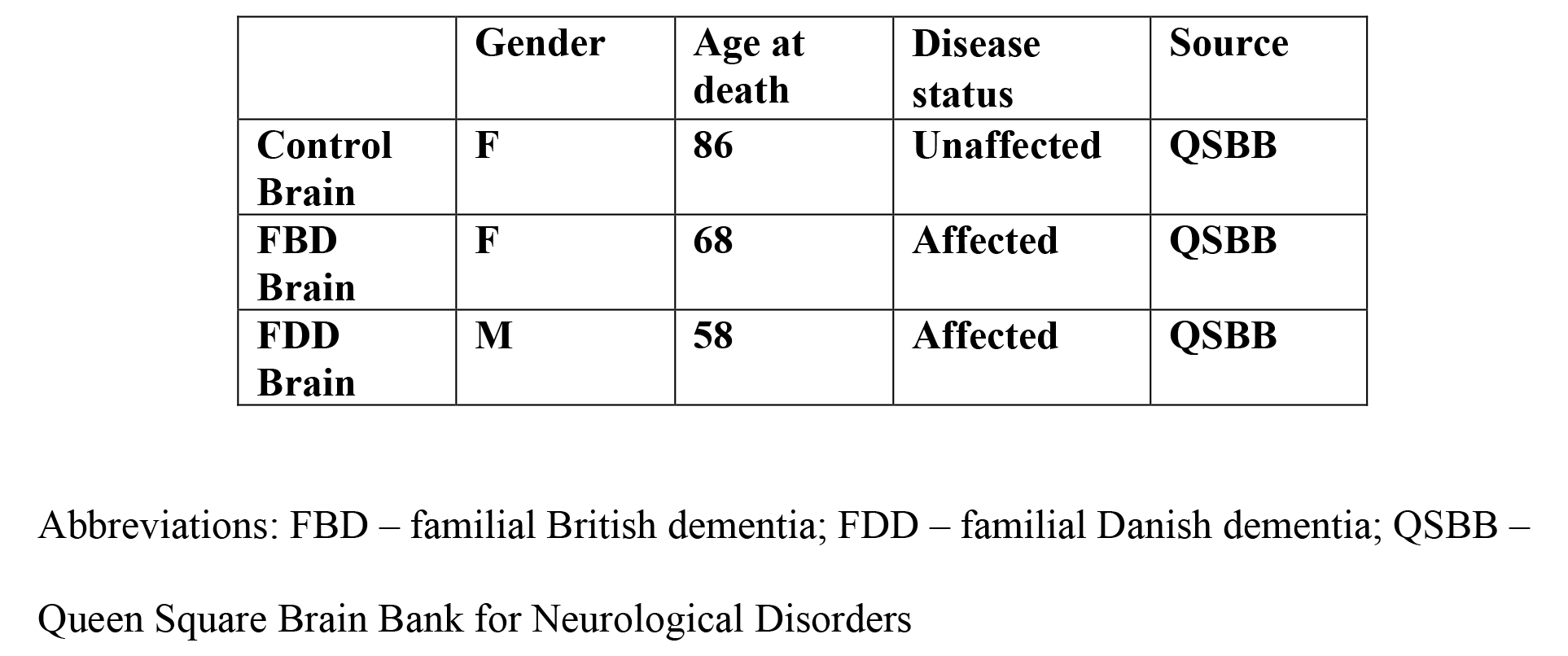
Details of tissue donors.

Finally, to further explore the functional relevance of *ITM2B/BRI2* expression in microglial activation in neurodegeneration, we performed gene coexpression network analysis to reveal the gene networks and the microglial states in which *ITM2B/BRI2* was expressed. Using a single cell RNA-sequencing dataset of microglia isolated from human Alzheimer’s disease brain and individuals with mild cognitive impairment (MCI) (Olah et al., 2020), we performed an improved version of weighted gene coexpression network analysis (WGCNA; (Botía et al., 2017)). The network analysis revealed a high enrichment of ARM signature genes within the *ITM2B/BRI2* network (Fig 5, Fig S5 and Supplementary Table 1). The genetic network containing *ITM2B/BRI2* contained known ARM genes such as *TREM2* and *TYROBP*, Complement-associated genes (*C1QA*, *C1QB*), lysosome-related genes (*CTSB*, *CTSS*, *LAMP2*) and *HLA* genes. To reinforce these findings, we leveraged additional online databases to further investigate coexpression networks for *ITM2B/BRI2* (see methods). Genes displaying coexpression with *ITM2B/BRI2* included the HLA gene *B2M* and lysosome-related genes *LAMP2*, *GRN, LAMP1*, *LAPTM4A*, *PSAP* and *ASAH1* (Fig S6). Similar to *ITM2B/BRI2*, the genes *B2M*, *ASAH1*, *GRN*, and *PSAP* were among the top 100 most highly expressed genes in iPSC-derived microglia (Supplementary Table 2) (Lin et al., 2018) and are also enriched in ARM states in additional human and mouse datasets (Keren-Shaul et al., 2017; Olah et al., 2020; Sala Frigerio et al., 2019). Together, these data support a normal, physiological role for *ITM2B/BRI2* in the microglial response to damage and disease, which is in addition to production of amyloidogenic peptides in FBD.

**Figure 5.**
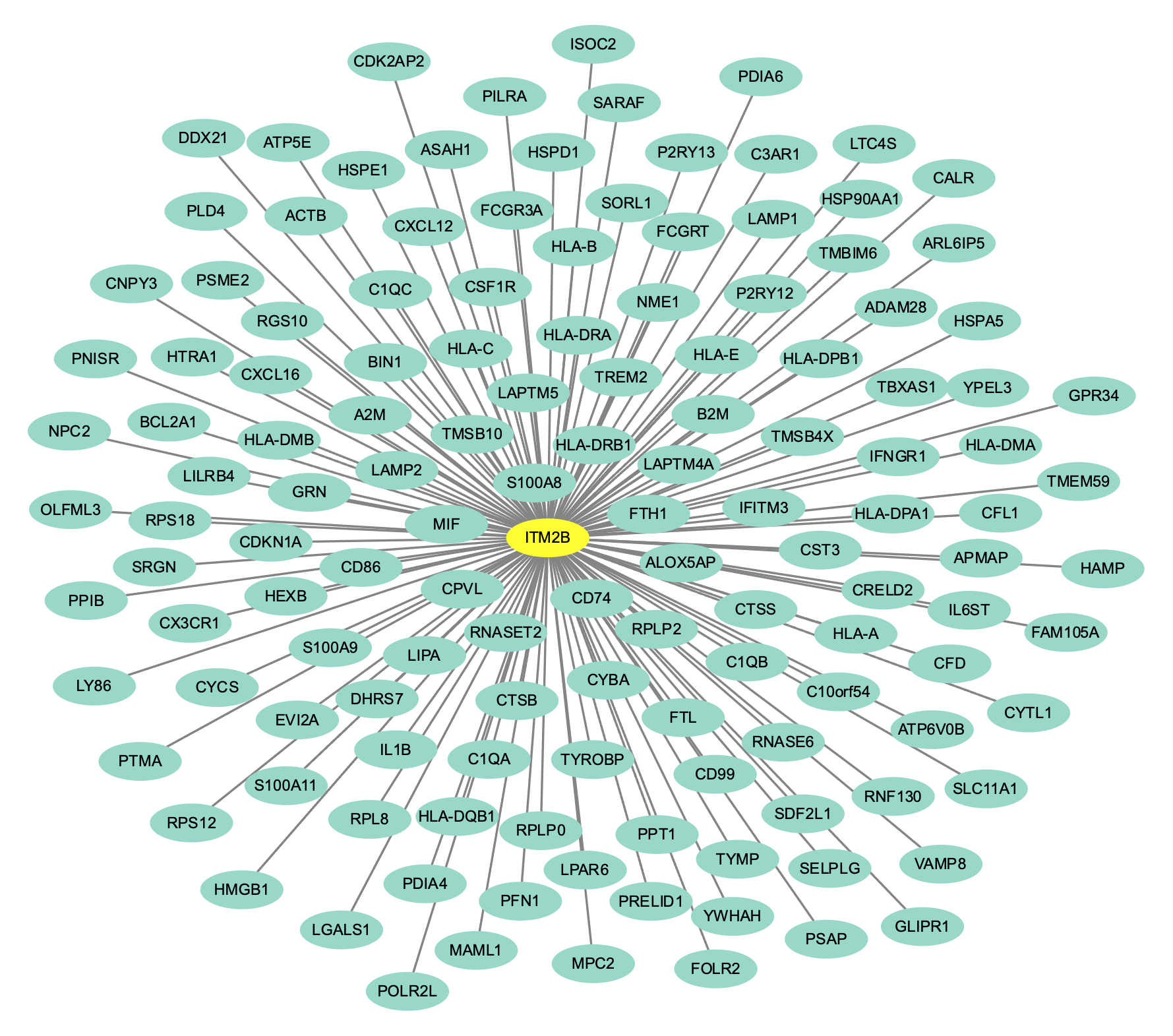
*ITM2B* is coexpressed with ARM genes in microglia isolated from human individuals showing neurodegeneration (Alzheimer’s disease and MCI). Genetic network plot of a module containing *ITM2B* detected in microglial cells isolated from human Alzheimer’s disease patients and individuals with MCI analysed by scRNA-seq (Olah et al., 2020). Genes with the highest connectivity to *ITM2B* from the co-expression network were plotted based on ranking the connectivity matrix of the expression data. This module contains genes associated with the DAM/ARM state (the full network and the strength of each interaction is given in Supplementary Table 1). Genes most strongly co-expressed with *ITM2B* include genes known to be associated with neurodegeneration including *LAPTM5*, HLA genes, *CTSB*, *CTSS*, *GRN*, *TREM2* and *TYROBP*. *ITM2B* is highlighted with a yellow oval.

## Discussion

Here we describe a novel patient-derived iPSC model of FBD, providing a human physiological model of disease. Surprisingly, expression of *ITM2B/BRI2* was substantially higher in microglia compared with neurons and astrocytes. Consequently, we were able to detect ABri in patient-derived microglial cultures and not in the neuronal cultures used in this study, suggesting microglia represent a major source of ABri in FBD. This is an unexpected finding and contrasts with some existing literature (Akiyama et al., 2004).

This finding supports the notion that in FBD, microglial-derived amyloidogenic peptides contribute to plaque pathology. Given the pathological and clinical overlap between FBD and Alzheimer’s disease, it is intriguing to consider the amyloid cascade hypothesis (Selkoe and Hardy, 2016) initiating via different amyloids from distinct cellular sources – converging on a disease pathogenesis featuring tau pathology, inflammation, neurodegeneration, and dementia-like symptoms.

Pathological examination of post-mortem tissue from FBD and FDD shows ABri/ADan colocalised with microglia in close proximity to pre-amyloid deposits. The presence of ABri and ADan in microglial cells in FBD and FDD respectively, highlights a critical role for microglia in either amyloid production or uptake. Further investigations would be needed to determine the exact role microglia play in conversion of amyloidogenic peptides to amyloid. Published bulk expression data suggests that *ITM2B/BRI2* expression is highest in the hippocampus and the cerebellum (Fig S7) (Hawrylycz et al., 2012; Ramasamy et al., 2014). This is distinct from the expression pattern another microglial marker *TREM2*. The expression levels correlate with the occurrence of parenchymal pathology in FBD (Holton et al., 2001) and FDD (Holton et al., 2002), whereas CAA is found more widespread. The levels of pathology in these brain structures may help to explain the clinical symptoms of disease.

Our data, together with existing expression data from human and mouse cells, show that *ITM2B/BRI2* expression is enriched in microglia (Friedman et al., 2018; Guttenplan et al., 2020; Hochgerner et al., 2018; Lake et al., 2018; Mancarci et al., 2017; Saunders et al., 2018; Zhang et al., 2016, 2014a). Indeed, we reanalysed data from Lin and colleagues who differentiated iPSC to neurons, astrocytes and microglia (Lin et al., 2018) and we saw that *ITM2B/BRI2* was within the top 100 highest expressed genes in the microglial lineage (based on normalised fragments per kilobase per million mapped fragments, Supplementary Table 2). Single cell data suggest that *ITM2B/BRI2* is enriched in ARM microglial clusters (Olah et al., 2020; Tuddenham et al., 2022). This leads to two potential hypotheses: 1) induction of a DAM/ARM-like state induces expression of *ITM2B/BRI2*, which leads to the production and deposition of ABri and further disease progression. Alternatively, 2) a putative loss of function of ITM2B/BRI2 protein, as described in mouse models (Tamayev et al., 2010; Yin et al., 2021), may negatively impact on the normal response of microglia to early pathological changes and cellular damage, thereby worsening the disease. We observed no evidence for reduced ITM2B/BRI2 protein abundance in our FBD patient-derived microglial model – supporting the first hypothesis or a combination of both.

Clinically, minor accidents and trauma have been associated with symptom onset in FBD, for example, a flu-like disease was associated with disease onset in one of the patients from whom iPSCs were made in this study (Harris et al., 2022). Whilst speculative, this might be compatible with a role for the immune system in FBD - whereby an inflammatory response to environmental factors may trigger or enhance expression of the pathological protein.

Despite the presence of ABri in iPSC-derived microglial cultures, we cannot discount the contribution of other cell types to ABri production; for example, endothelial cells and oligodendrocytes; especially given the high burden of angiopathy (Holton et al., 2001). However, in situ hybridisation for *ITM2B/BRI2* in post-mortem tissue displayed weak signal in white matter and the vascular unit (Lashley et al., 2008). Additionally, neurons have been shown to express *ITM2B/BRI2* (Akiyama et al., 2004) and may upregulate ITM2B in a context dependent manner (Chen et al., 2020). Indeed, our iPSC-derived neuronal model does demonstrate low level expression. However, single cell sequencing datasets (Fig S1 and S2) and the data presented herein support the finding that microglia are a major contributing source of ITM2B/BRI2 and ABri.

In summary, we propose a central role for microglial-derived ABri in FBD and subsequent non-cell autonomous mechanisms driving neuronal dysfunction. This surprising finding has relevance to the amyloid cascade hypothesis and Alzheimer’s disease; insomuch as 1) distinct origins of amyloidogenic peptides can culminate in neurodegeneration and dementia-like symptoms and 2) microglia and the immune response are central to disease onset as well as progression.

## Materials and Methods

### Cell Culture

Patient-derived fibroblasts were obtained from a skin biopsy with ethical approval from the Institute of Neurology joint research ethics committee at the Hospital for Neurology and Neurosurgery (10/H0721/87) with informed consent (Table 1). Fibroblasts were grown in DMEM supplemented with 10% FBS and passaged using 0.05% trypsin. Fibroblasts, below passage 4, were reprogrammed using episomal reprogramming as described previously (Okita et al., 2011). Episomal plasmids, obtained from Addgene #27077, #27078 and #27080, were electroporated into fibroblasts using Lonza P2 Nucleofection. 7 days post electroporation, media was changed to Essential 8 and iPSC colonies appeared after a subsequent 20 days. iPSC colonies were picked manually and expanded in Essential 8 media, on geltrex substrate and passaged manually using 0.5mM EDTA. Ctrl1 and Ctrl2 refer to the well characterised RBi001-a and SIGi1001-a-1, respectively, both available via Sigma Aldrich (Arber et al., 2021, 2020).

Genomic DNA was isolated from iPSC clones using cell lysis buffer containing 0.5% SDS and 0.5mg/ml proteinase k. Samples were lysed at 55°C overnight and DNA was extracted using phenol-chloroform extraction with ethanol-based precipitation. DNA was quantified using nanodrop, diluted to 50 ng/µl in 15 µl and put forward to Sanger sequencing using standard PCR master mix in a touchdown-PCR.

Karyotype stability was confirmed by The Doctors Laboratory (London, UK) using G-band analysis. Stem cell phenotype was confirmed via comparing the expression of 770 genes associated with pluripotency and differentiation with a panel of 3 established iPSC/hESC (Ctrl1, Ctrl2 and Shef6 (Arber et al., 2020)) lines using the Nanostring Stem Cell Characterisation Panel.

iPSCs were differentiated to cortical neurons using established protocols (Arber et al., 2020; Shi et al., 2012). Briefly, iPSCs at 100% confluence were subject to neural induction using dual SMAD inhibition in N2B27 media (1μM dorsomorphin and 10μM SB431542, both TOCRIS). Day 90 was taken as the final timepoint.

iPSCs were differentiated to astrocytes following established protocols (Hall et al., 2017). Neuronal cultures (as above) were enriched for astrocytes via continuous EDTA passaging in N2B27 media containing 10ng/ml FGF2 (Peprotech). At 150 days in vitro, a final two-week maturation step involved BMP4 (10ng/ml) and LIF (10ng/ml) in N2B27 media.

iPSCs were differentiated to microglia following established methods (Garcia-Reitboeck et al., 2018; Xiang et al., 2018). Briefly, myeloid embryoid bodies (EBs) were produced using 10,000 cells in Essential 8 media supplemented with 50ng/ml BMP4, 50ng/ml VEGF and 20ng/ml SCF. After 4 days myeloid EBs were maintained in X-Vivo 15 media supplemented with 100ng/ml MCSF and 25ng/ml IL3. Microglia-like cells were harvested and matured using DMEM-F12 media supplemented with 100ng/ml IL34, 25ng/ml MCSF and 5ng/ml TGFβ1. A final maturation was performed via a 2 day treatment with CX3CL1 (100ng/ml) and CD200 (100ng/ml).

All components were ThermoFisher, unless stated. All growth factors were Peprotech, unless stated.

### Participant details

### qPCR

RNA was extracted from cells using Trizol, following the manufacturers protocol. 2µg of RNA was reverse transcribed using SuperScript IV reverse transcriptase, random hexamer mix and RNAse OUT. qPCR was run on an Agilent Aria MX using POWER Sybr green master mix. Primers are shown in Table 3.

**Table 3.**
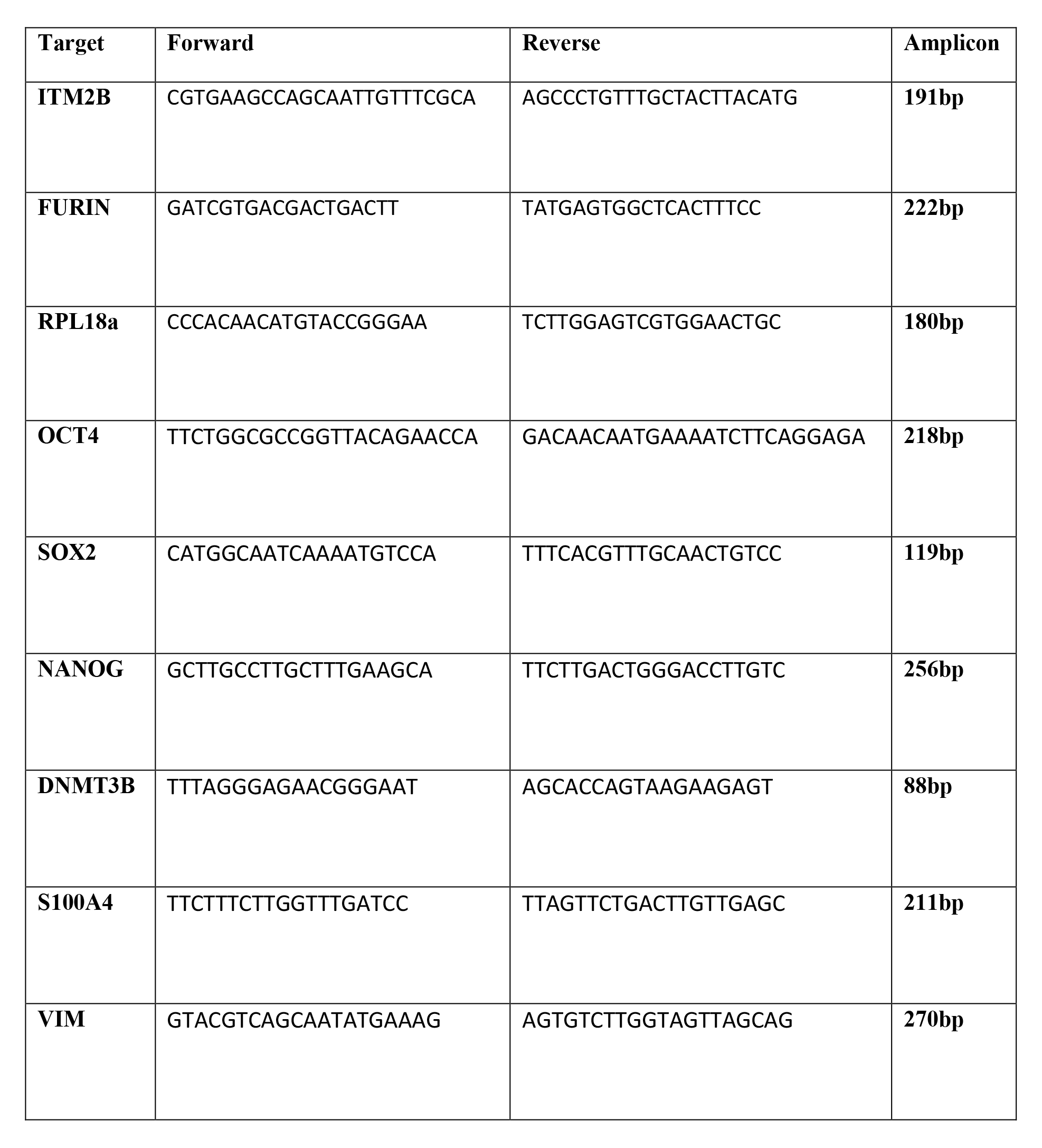
Primers used in this study

### Immunocytochemistry

Cells were fixed in 4% paraformaldehyde for 15 minutes. Cells were then washed thrice in 0.3% Triton-X-100 in PBS (PBSTx) prior to blocking in 3% bovine serum albumin in PBST. Primary antibodies (Table 4) were incubated in blocking solution overnight. After three subsequent washes in PBSTx, secondary antibodies (AlexFluor) were incubated for 1 hour in the dark in blocking solution. After a final three washes and exposure to DAPI nuclear stain, images were taken on a Zeiss LSM microscope. No post-hoc image adjustments were made.

**Table 4.** Antibodies used in this study.

| **Target** | **Species** | **Company** | **RRID** |
| --- | --- | --- | --- |
| SSEA4 | mouse | BioLegend<br>330401 | AB_1089209 |
| OCT4 | goat | Santa Cruz<br>sc-5297 | AB_628051 |
| TBR1 | rabbit | Abcam | AB_2200219 |
|  |  | ab31940 |  |
| TUJ1 | mouse | BioLegend<br>801201 | AB_2313773 |
| IBA1 | goat | Abcam<br>ab107159 | AB_10972670 |
| ITM2B/BRI2<br>CTF | rabbit | Sigma<br>HPA029292 | AB_10601917 |
| ITM2B/BRI2<br>NTF | mouse | Santa cruz<br>sc-374362 | AB_10988049 |
| TREM2 | rabbit | Cell Signaling<br>Tech<br>91068 | AB_2721119 |
| SOX9 | Rabbit | Abcam<br>ab185966 | AB_2728660 |
| Actin | mouse | Sigma<br>A1978 | AB_476692 |
| GAPDH | mouse | Ambion<br>AM4300 | AB_437392 |
| ABri | rabbit | 338 (Vidal et<br>al., 1999) |  |
| ADan | rabbit | 5282 (Vidal et<br>al., 2000) |  |
| CD68 | mouse | Dako M0876 | AB_2074844 |
| CR3/43 | mouse | Dako M0775 | AB_2313661 |

### Immunohistochemistry

Formalin fixed paraffin embedded sections were deparaffinized in xylene, followed by rehydration using graded alcohols (100%, 95% and 70%). For all immunohistochemical staining the endogenous peroxidase activity was blocked using 0.3% H_2_O_2_ in methanol for 10 minutes. Sections were subjected to various pre-treatments, depending on the antibody used. For ABri and ADan, sections were pre-treated with formic acid for 10 minutes. For microglial staining (CD68 or CR3/43), sections were pressure cooked in citrate buffer pH6.0 for 10 mins. Non-specific protein binding was blocked using 10% non-fat milk in PBS (0.05M pH7.2) by incubating the sections for 30 minutes at room temperature. Sections were incubated with the required primary antibody (ABri: 338 1:1000 from Ghiso lab; ADan: 5282 1:1000 from Ghiso lab; CD68: 1:150 DAKO; CR3/43: 1:100 DAKO) for 1 hour at room temperature. Incubation with the relevant biotinylated secondary antibody (Vector) was carried out for 30 minutes at room temperature. Sections were incubated in avidin-biotin complex (ABC, Dako) for 30 minutes and the antigen-antibody reaction was visualized using di-aminobenzidine (DAB, Sigma) as the chromogen. Sections were counterstained with Mayers haematoxylin (BDH), dehydrated and mounted.

### Thioflavin staining

Thioflavin staining was used to demonstrate protein deposits in amyloid conformation and used in addition to immunohistochemical staining. Once immunohistochemical staining was complete sections were incubated with Thioflavin solution (0.1% aqueous solution) for 7 minutes and differentiated with 70% alcohol, followed by washing in distilled water.

### Western Blotting

Cells were lysed in RIPA buffer with protease and phosphatase inhibitors (Roche) and centrifuged to remove insoluble debris. Protein content was quantified via BCA assay (BioRad). Samples were denatured at 95°C for 5 minutes in LDS buffer with DTT and loaded on 4-12% precast polyacrylamide gels. Gels were transferred to nitrocellulose membranes and blocked, using PBS with 0.1% Tween-20 (PBSTw) and 3% BSA. Primary antibodies were incubated in blocking solution overnight, washed thrice in PBSTw and incubated with secondary antibodies for 1 hour. After three final washes, images were captured on a LiCor Odyssey fluorescent imager.

### Gene CoExpression Analysis

We employed online databases on human tissue including HumanBase (human macrophages, top 20 genes, minimum interaction prediction confidence: 0.65), GeneFriends (human brain tissue, top 10 genes, Pearson correlation threshold: 0.85) and COXPRESdb v8 (non-specific tissue, top 10 co-expressed genes ranked based on z-scores) to generate co-expression networks for *ITM2B* (Table 5). These lists were analysed to determine the genes which were present in multiple databases.

**Table 5:** Online Platforms for Co-expression Network Analysis

| Platform |  | Description |
| --- | --- | --- |
| HumanBase 1.0<br>(formerly<br>GIANT)<br>(Greene et al.,<br>2015) | https://humanbase.flatironinstitute.org/gene/ | A resource that includes genome-scale functional maps and tissue-specific networks across 144 human tissues with statistical prioritisation developed using Bayesian framework on more than 14000 publications. Human macrophage tissue was selected for this analysis. |
| GeneFriends<br>(Raina et al.,<br>2023) | https://www.genefriends.org/ | A guilt-by-association data-driven analysis platform using RNAseq based gene co-expression network construction with functional annotation for multiple species. Human brain tissue was selected for this analysis. |
| COXPRESdb<br>v8(Obayashi et<br>al., 2023) | https://coxpresdb.jp/ | A gene coexpression database based on DNA microarray analysis and over 200,000 RNASeq runs which collates data to present coexpression |
|  |  | relationships based on relative expression patterns of genes and networks and protein-protein-interactions. This platform also uses the KEGG pathway and Gene Ontology Biological Process scores to substantiate the results. |

The ITM2B gene network (Fig 5) was generated through a co-expression analysis of pre-processed Pearson’s residuals obtained from a microglial single cell RNA-seq dataset collected from the dorsolateral prefrontal cortex of Alzheimer’s Disease (AD) brains (Olah et al., 2020). The dataset was filtered to remove cells that appeared unhealthy or potential doublets, with cells having greater than 5% ratio of mitochondrial to total counts or less than 1000 counts or less than 700 genes detected being removed using Seurat’s subset function. Additionally, samples from individuals with epilepsy were removed from the analysis, as they are not fully representative of healthy brain. Genes showing low variation in expression between cells (coefficient of variation for Pearson’s residuals <15%) were also removed, as they are not informative for co-expression analysis. The co-expression analysis was performed using the "getDownstreamNetwork" function from CoExpNets, which is an optimized version of the popular weighted gene co-expression network analysis (WGCNA) package (Botía et al., 2017). This optimization involves a k-means clustering step to re-categorize genes into biologically relevant and reproducible modules. The mean log2 normalized expression of the most central genes within the "turquoise" module was used to determine *ITM2B* as one of the hub genes in the network. These genes were ranked based on their module membership (MM) scores, which were calculated using the "getMM" function from CoExpNets. The correlation matrix of the expression data was then used to rank the most connected genes to *ITM2B* within the module. The resulting network was visualized using the Cytoscape v.3.9.1 software.

### Gene knockdown

*ITM2B/BRI2* was knocked down in iPSC-derived microglial-like cells using DharmaFECT 1 (Horizon) using SMARTpool siRNA (Horizon) alongside scrambled, non-targeting siRNA. Briefly, cells were plated at 500,000 cells per well of a 6 well plate. siRNA was prepared in DharmaFECT 1 (2.5µl per well) in serum free media for 20 minutes, before evenly distributing onto cells. Protein lysates were taken 72 hours later and run for Western blots, as above.

### Funding

CA was supported by an Alzheimer’s Society fellowship (AS-JF-18-008). JMC was supported by an EPSRC PhD studentship (EP/N509577/1). DAS, JH, NR, and UY research was funded by the UK DRI, which receives its funding from the DRI Ltd, funded by the UK Medical Research Council, Alzheimer’s Society and ARUK. JH is supported by the Dolby Foundation. SW is supported by senior research fellowship from Alzheimer’s Research UK (ARUK-SRF2016B-2). TMP was supported by funding to JMP and J Hardy from the Innovative Medicines Initiative 2 Joint Undertaking under grant agreement No 115976. This Joint Undertaking receives support from the European Union’s Horizon 2020 research and innovation programme and EFPIA. This work was supported by the Medical Research Council (MR/M02492X/1), the National Institute of Health (RF1 AG059695) and by the National Institute for Health and Care Research University College London Hospitals Biomedical Research Centre.

---

**Supplementary Table 1. Weighted gene coexpression network analysis of *ITM2B* in human microglia.** Connectivity between genes with *ITM2B* displayed in Figure 5, representing the strength of interaction.

**Supplementary Table 2. Top 100 expressed genes in iPSC-derived Microglia based on FPKM from Lin et al. (2018).**

**Figure S1.**
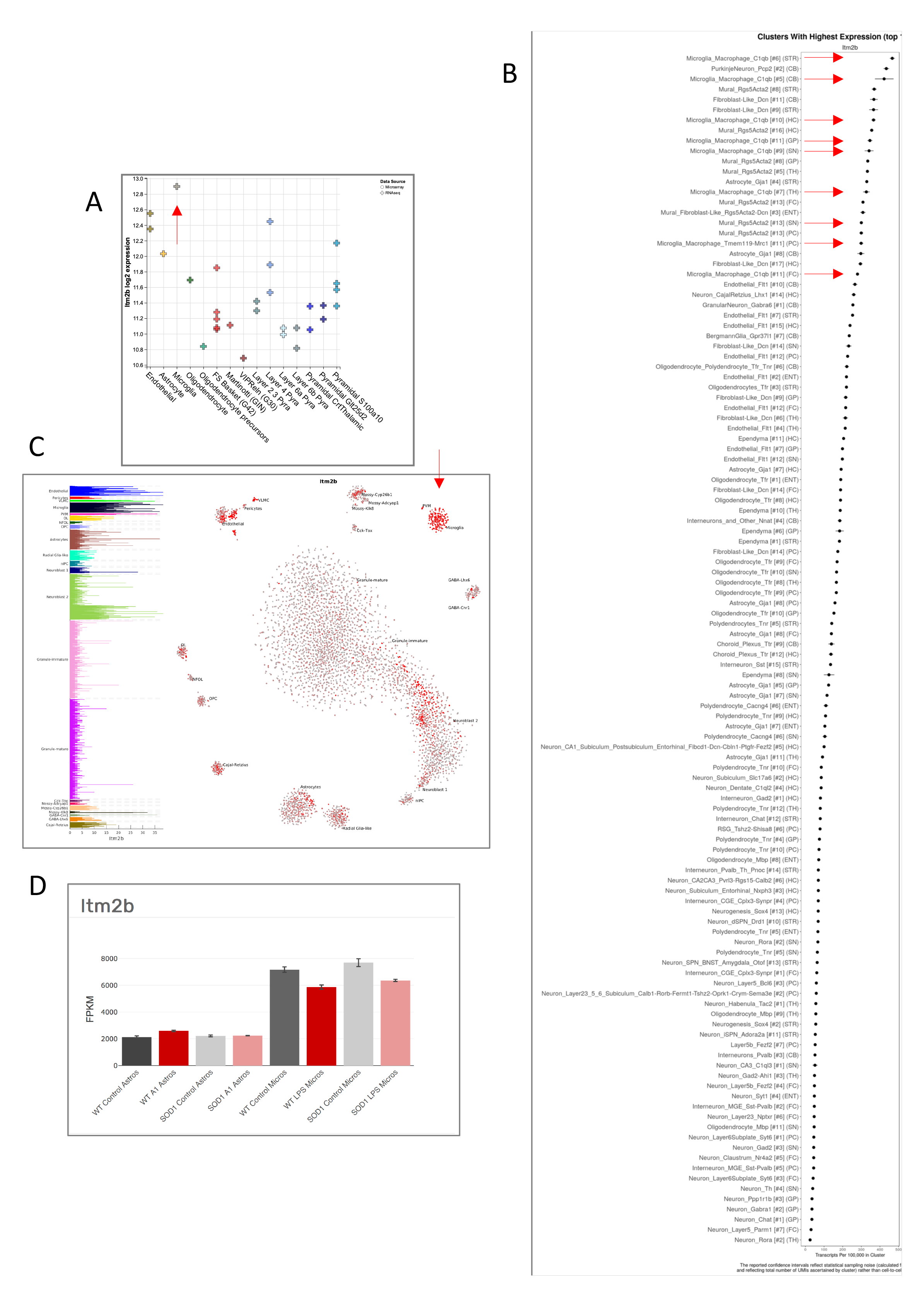
Data of *Itm2b* expression in different mouse brain cell types from published studies. Red arrows highlight microglial datapoints. A) Data from NeuroExpresso (Mancarci et al., 2017), B) Data from DropVis (Saunders et al., 2018), C) data from (Hochgerner et al., 2018), D) data from GliaSeq (Guttenplan et al., 2020).

**Figure S2.**
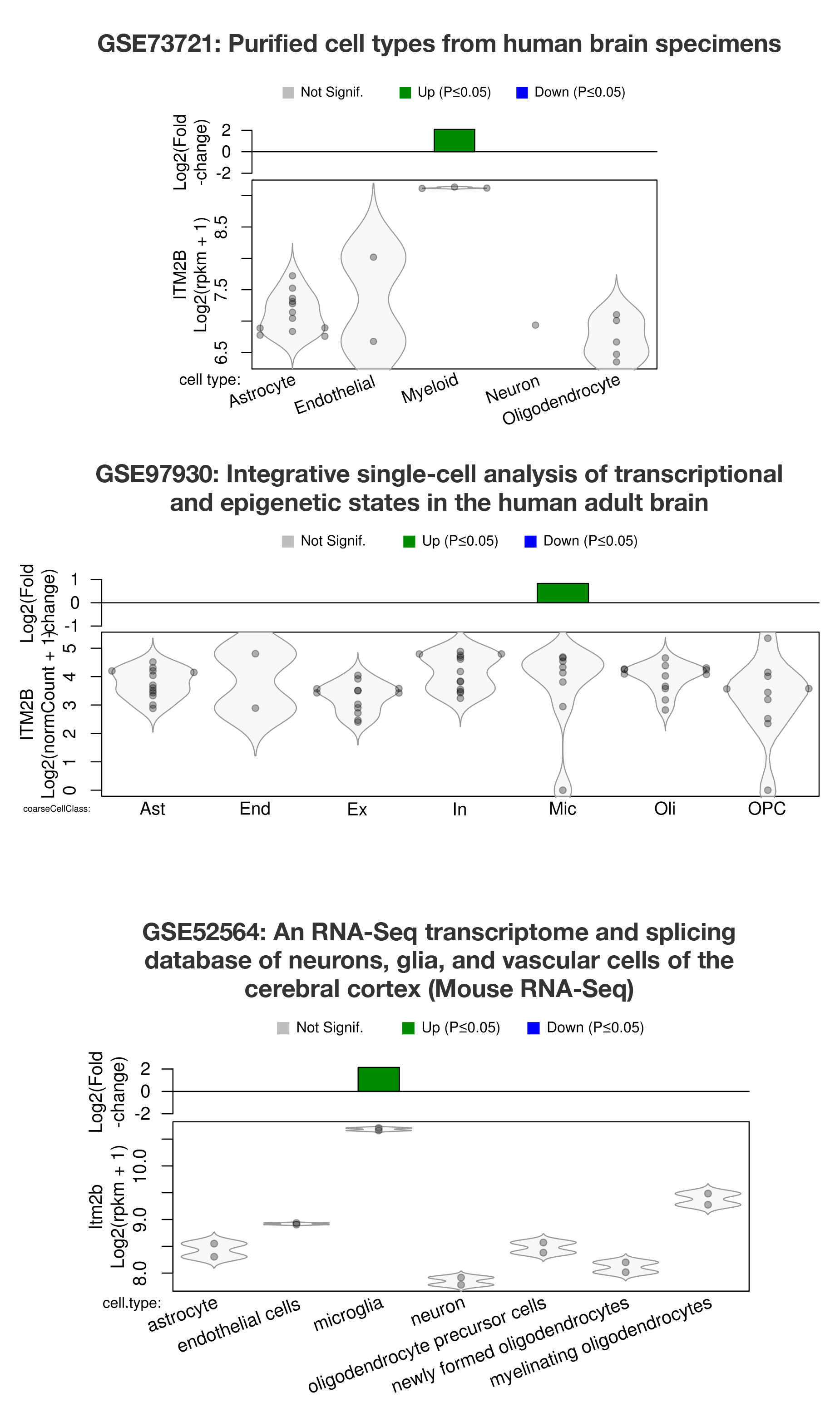
**Preferential expression of *ITM2B* and orthologue *Itm2b* in microglia versus other cell types in human and mouse brain**. Top panel: data from purified cell types of human brain, GSE73721 (Zhang et al., 2016). Middle panel: data from human brain analysed by single nucleus sequencing, GSE97930 (Lake et al., 2018). Bottom panel: data from purified cell types of mouse cerebral cortex, GSE52564 (Zhang et al., 2014b). Data was quantified and visualised from the Myeloid Landscape portal (http://research-pub.gene.com/BrainMyeloidLandscape; (Friedman et al., 2018).

**Figure S3.**
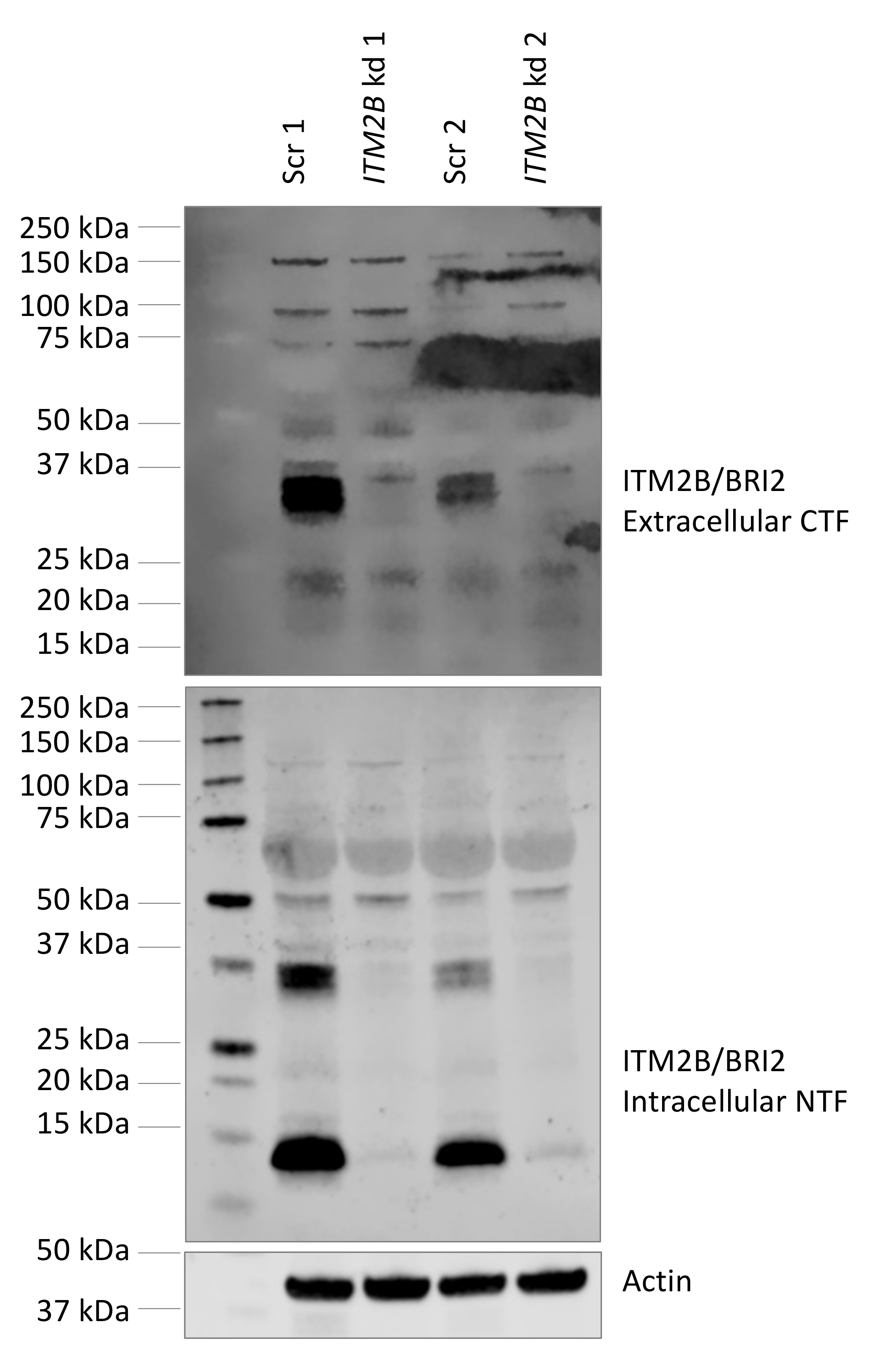
Specificity of ITM2B antibody. iPSC-microglia were treated with scrambled (scr) or (*ITM2B*) siRNA (kd) (SMART pool, Dharmacon) and protein was isolated 72 hours later. Knockdown experiments from two independent control cell lines are shown. Western blotting confirms the specificity of antibodies HPA029292 and sc-374362.

**Figure S4.**
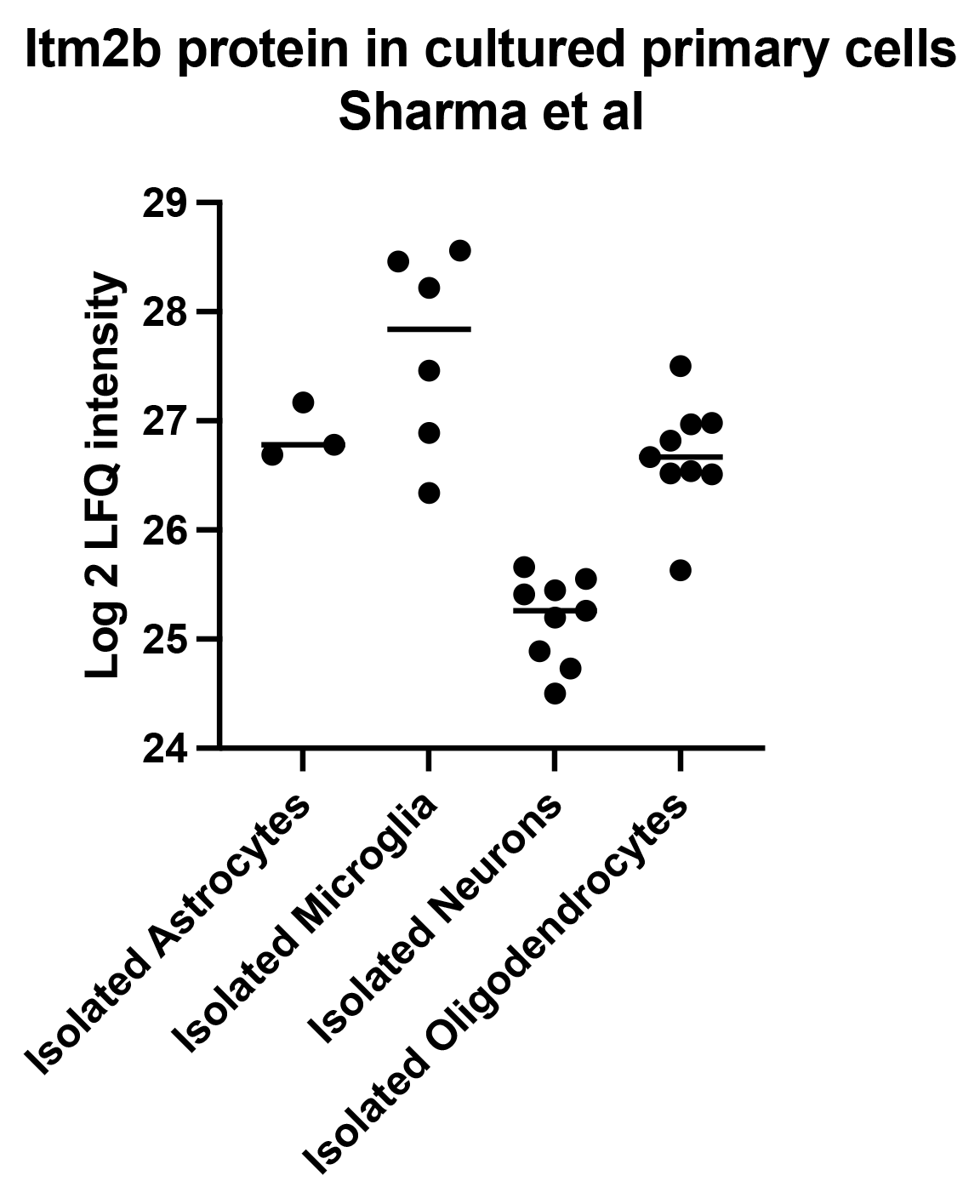
Proteomic data on sorted and cultured mouse brain cell types. Isolated mouse microglia have higher expression of ITM2B at the protein level in published proteomic data (Sharma et al., 2015).

**Figure S5.**
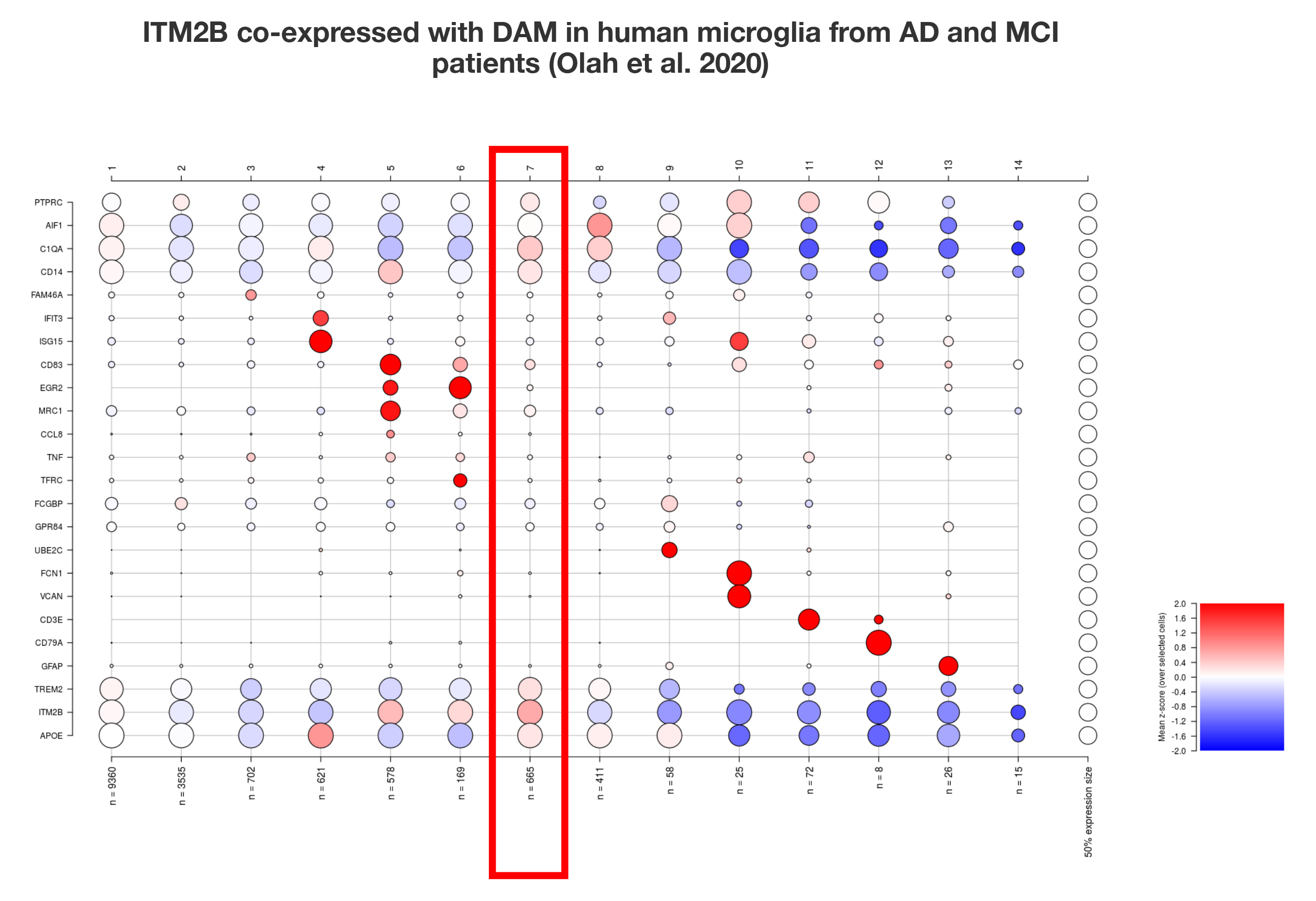
*ITM2B* is co-expressed with ARM genes in human microglia. Single cell expression data from human microglia, distinguished into 14 clusters (Olah et al., 2020). *ITM2B* is enriched in module 7 (red box) (circle size relates to fraction of cells expressing the gene and colour relates to z-score). This module contains *TREM2*, *APOE* and *C1QA* and was described to have the strongest enrichment for ARM genes.

**Figure S6.**
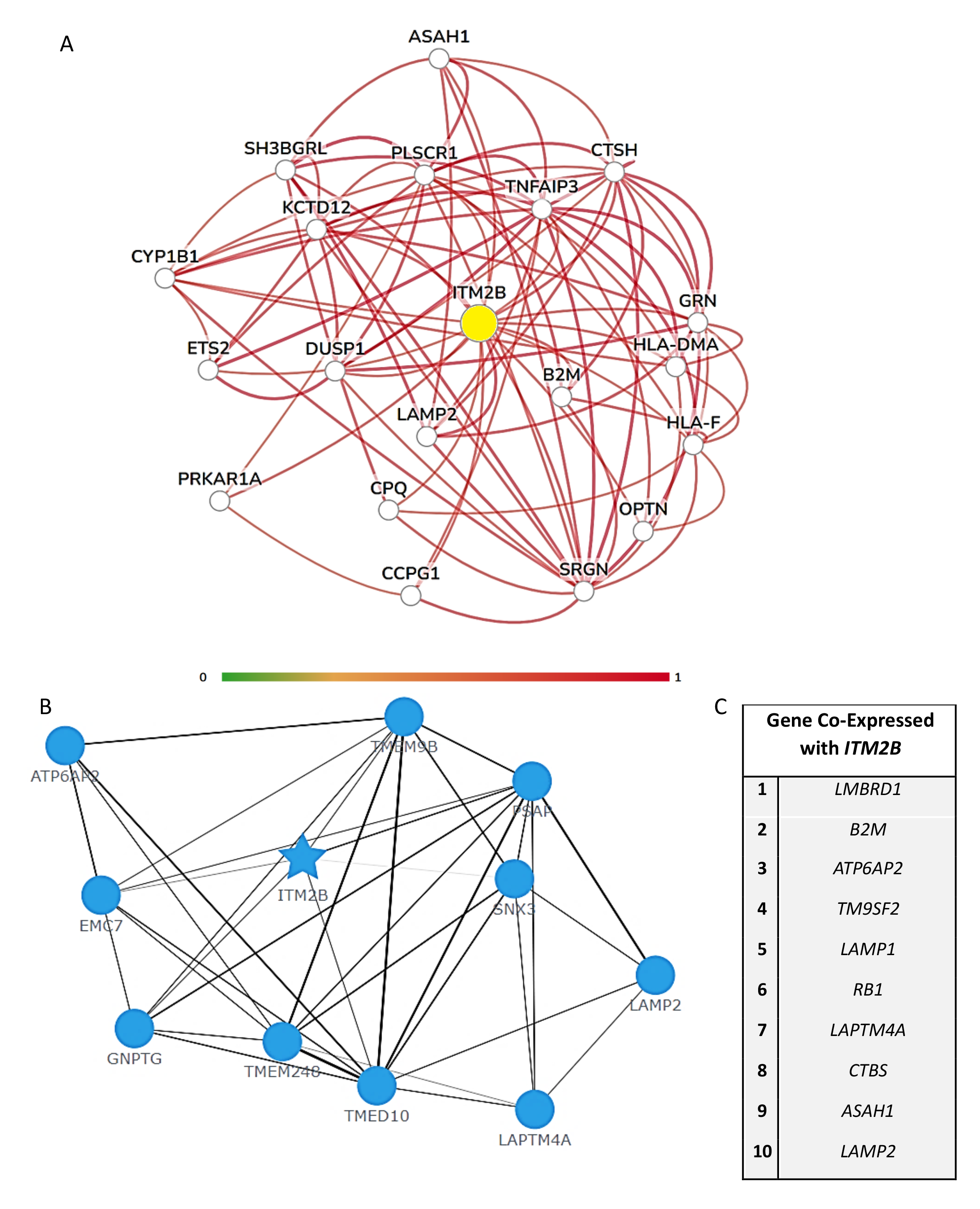
*ITM2B* Co-expression Networks. Online databases including HumanBase on human macrophages (A), GeneFriends on human brain tissue (B) and COXPRESdb human non-specific tissue (C) were analysed according to minimum interaction prediction confidence 0.65, Pearson correlation threshold 0.85 and z-score ranking respectively.

**Figure S7.**
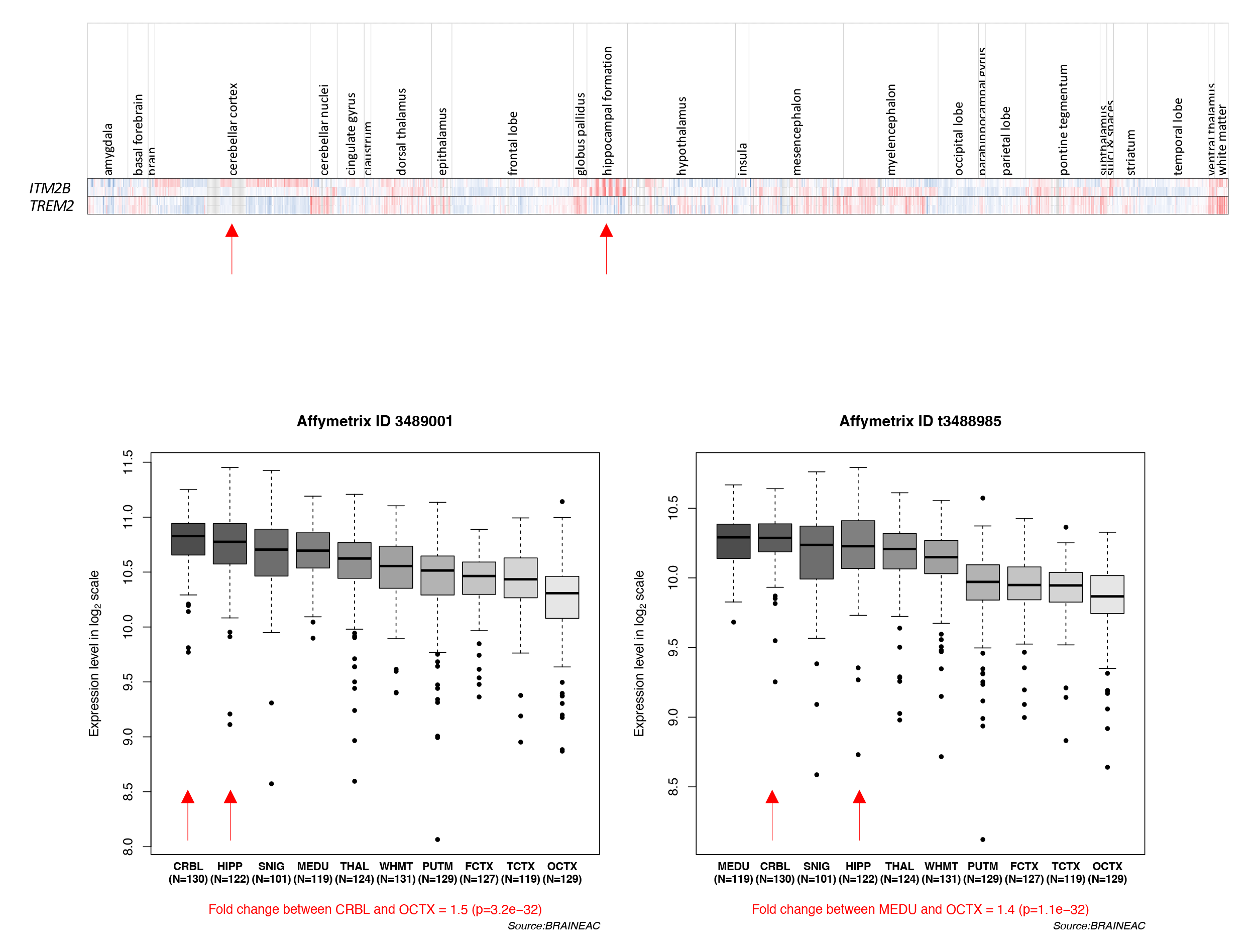
*ITM2B* expression in different brain regions. A) Expression of *ITM2B/BRI2* across brain regions in comparison to the microglial gene *TREM2*. Data from Allan Human Brain Atlas available from human.brain-map.org RRID:SCR_007416 (Hawrylycz et al., 2012). B) Expression of *ITM2B/BRI2* in different brain regions from braineac.org. 2 representative transcripts of 11 are shown (Ramasamy et al., 2014).

