## Supplementary Tables 1 and 2 for "Microglia produce the amyloidogenic ABri peptide in familial British dementia"

Supplementary Table 1

| | from | to | Zero | MOO39 | value |
|---|---|---|---|---|---|
| 1 | ITM2B | ITM2B | 0 | M0039 | 1 |
| 2 | ITM2B | RNASET2 | 0 | M0039 | 0.00111991 |
| 3 | ITM2B | GPR34 | 0 | M0039 | 0.00031787 |
| 4 | ITM2B | HLA-DRB1 | 0 | M0039 | 0.00029632 |
| 5 | ITM2B | CPVL | 0 | M0039 | 0.00022225 |
| 6 | ITM2B | HLA-DPA1 | 0 | M0039 | 0.00019663 |
| 7 | ITM2B | ASAH1 | 0 | M0039 | 0.00017145 |
| 8 | ITM2B | CD74 | 0 | M0039 | 0.00014948 |
| 9 | ITM2B | TMSB10 | 0 | M0039 | 0.00014778 |
| 10 | ITM2B | HLA-DRA | 0 | M0039 | 0.0001403 |
| 11 | ITM2B | FOLR2 | 0 | M0039 | 0.00014021 |
| 12 | ITM2B | CALR | 0 | M0039 | 0.00013646 |
| 13 | ITM2B | HSP90B1 | 0 | M0039 | 0.00012706 |
| 14 | ITM2B | IFNGR1 | 0 | M0039 | 0.00011757 |
| 15 | ITM2B | IFITM3 | 0 | M0039 | 0.00011703 |
| 16 | ITM2B | HSPA5 | 0 | M0039 | 9.80E-05 |
| 17 | ITM2B | FCGRT | 0 | M0039 | 8.32E-05 |
| 18 | ITM2B | SDF2L1 | 0 | M0039 | 8.00E-05 |
| 19 | ITM2B | TUBA1B | 0 | M0039 | 7.53E-05 |
| 20 | ITM2B | LAPTM5 | 0 | M0039 | 7.47E-05 |
| 21 | ITM2B | ALOX5AP | 0 | M0039 | 7.30E-05 |
| 22 | ITM2B | LAPTM4A | 0 | M0039 | 6.82E-05 |
| 23 | ITM2B | C10orf54 | 0 | M0039 | 5.99E-05 |
| 24 | ITM2B | HTRA1 | 0 | M0039 | 5.97E-05 |
| 25 | ITM2B | RNASE6 | 0 | M0039 | 5.82E-05 |
| 26 | ITM2B | SELPLG | 0 | M0039 | 5.81E-05 |
| 27 | ITM2B | ARL6IP5 | 0 | M0039 | 4.99E-05 |
| 28 | ITM2B | PILRA | 0 | M0039 | 4.82E-05 |
| 29 | ITM2B | PSAP | 0 | M0039 | 4.53E-05 |
| 30 | ITM2B | RNF130 | 0 | M0039 | 4.24E-05 |
| 31 | ITM2B | HLA-B | 0 | M0039 | 4.23E-05 |
| 32 | ITM2B | HLA-DMB | 0 | M0039 | 3.94E-05 |
| 33 | ITM2B | FAM105A | 0 | M0039 | 3.77E-05 |
| 34 | ITM2B | A2M | 0 | M0039 | 3.69E-05 |
| 35 | ITM2B | CTSB | 0 | M0039 | 3.68E-05 |
| 36 | ITM2B | CST3 | 0 | M0039 | 3.57E-05 |
| 37 | ITM2B | P2RY12 | 0 | M0039 | 3.48E-05 |
| 38 | ITM2B | HLA-C | 0 | M0039 | 3.47E-05 |
| 39 | ITM2B | B2M | 0 | M0039 | 3.38E-05 |
| 40 | ITM2B | HLA-A | 0 | M0039 | 2.94E-05 |
| 41 | ITM2B | P2RY13 | 0 | M0039 | 2.91E-05 |
| 42 | ITM2B | S100A8 | 0 | M0039 | 2.78E-05 |
| 43 | ITM2B | TMEM59 | 0 | M0039 | 2.64E-05 |
| 44 | ITM2B | TYMP | 0 | M0039 | 2.63E-05 |
| 45 | ITM2B | SARAF | 0 | M0039 | 2.52E-05 |

| | | | | | |
|---|---|---|---|---|---|
| **46** | *ITM2B* | *PLD4* | 0 | M0039 | 2.50E-05 |
| **47** | *ITM2B* | *APMAP* | 0 | M0039 | 2.39E-05 |
| **48** | *ITM2B* | *FTL* | 0 | M0039 | 2.30E-05 |
| **49** | *ITM2B* | *CSF1R* | 0 | M0039 | 2.27E-05 |
| **50** | *ITM2B* | *C3AR1* | 0 | M0039 | 2.23E-05 |
| **51** | *ITM2B* | *CFD* | 0 | M0039 | 2.17E-05 |
| **52** | *ITM2B* | *ATP6V0B* | 0 | M0039 | 2.17E-05 |
| **53** | *ITM2B* | *DHRS7* | 0 | M0039 | 2.01E-05 |
| **54** | *ITM2B* | *CX3CR1* | 0 | M0039 | 1.93E-05 |
| **55** | *ITM2B* | *HAMP* | 0 | M0039 | 1.89E-05 |
| **56** | *ITM2B* | *CXCL16* | 0 | M0039 | 1.88E-05 |
| **57** | *ITM2B* | *HLA-DPB1* | 0 | M0039 | 1.78E-05 |
| **58** | *ITM2B* | *CYBA* | 0 | M0039 | 1.77E-05 |
| **59** | *ITM2B* | *HLA-DMA* | 0 | M0039 | 1.75E-05 |
| **60** | *ITM2B* | *CTSS* | 0 | M0039 | 1.69E-05 |
| **61** | *ITM2B* | *GLIPR1* | 0 | M0039 | 1.68E-05 |
| **62** | *ITM2B* | *PDIA4* | 0 | M0039 | 1.66E-05 |
| **63** | *ITM2B* | *S100A9* | 0 | M0039 | 1.63E-05 |
| **64** | *ITM2B* | *SORL1* | 0 | M0039 | 1.53E-05 |
| **65** | *ITM2B* | *S100A11* | 0 | M0039 | 1.49E-05 |
| **66** | *ITM2B* | *MANF* | 0 | M0039 | 1.40E-05 |
| **67** | *ITM2B* | *ATP5E* | 0 | M0039 | 1.38E-05 |
| **68** | *ITM2B* | *HSP90AA1* | 0 | M0039 | 1.20E-05 |
| **69** | *ITM2B* | *BCL2A1* | 0 | M0039 | 1.12E-05 |
| **70** | *ITM2B* | *NME1* | 0 | M0039 | 1.10E-05 |
| **71** | *ITM2B* | *PTMA* | 0 | M0039 | 1.08E-05 |
| **72** | *ITM2B* | *RPLP2* | 0 | M0039 | 1.06E-05 |
| **73** | *ITM2B* | *CD99* | 0 | M0039 | 1.05E-05 |
| **74** | *ITM2B* | *GRN* | 0 | M0039 | 1.03E-05 |
| **75** | *ITM2B* | *FCGR3A* | 0 | M0039 | 1.01E-05 |
| **76** | *ITM2B* | *CD86* | 0 | M0039 | 9.41E-06 |
| **77** | *ITM2B* | *LILRB4* | 0 | M0039 | 9.35E-06 |
| **78** | *ITM2B* | *PPIB* | 0 | M0039 | 9.07E-06 |
| **79** | *ITM2B* | *FTH1* | 0 | M0039 | 8.83E-06 |
| **80** | *ITM2B* | *TMSB4X* | 0 | M0039 | 8.79E-06 |
| **81** | *ITM2B* | *RGS10* | 0 | M0039 | 8.06E-06 |
| **82** | *ITM2B* | *LIPA* | 0 | M0039 | 7.99E-06 |
| **83** | *ITM2B* | *HSPE1* | 0 | M0039 | 7.96E-06 |
| **84** | *ITM2B* | *CFL1* | 0 | M0039 | 7.70E-06 |
| **85** | *ITM2B* | *DBI* | 0 | M0039 | 7.67E-06 |
| **86** | *ITM2B* | *BHLHE41* | 0 | M0039 | 7.64E-06 |
| **87** | *ITM2B* | *CYTL1* | 0 | M0039 | 7.30E-06 |
| **88** | *ITM2B* | *SERF2* | 0 | M0039 | 7.28E-06 |
| **89** | *ITM2B* | *MIF* | 0 | M0039 | 7.26E-06 |
| **90** | *ITM2B* | *ADAM28* | 0 | M0039 | 7.04E-06 |
| **91** | *ITM2B* | *PPT1* | 0 | M0039 | 6.75E-06 |
| **92** | *ITM2B* | *LPAR6* | 0 | M0039 | 6.60E-06 |
| **93** | *ITM2B* | *YPEL3* | 0 | M0039 | 6.59E-06 |

| | | | | | |
|---|---|---|---|---|---|
| **94** | *ITM2B* | *CRELD2* | 0 | M0039 | 6.33E-06 |
| **95** | *ITM2B* | *PFN1* | 0 | M0039 | 6.30E-06 |
| **96** | *ITM2B* | *RPS18* | 0 | M0039 | 6.15E-06 |
| **97** | *ITM2B* | *RPS12* | 0 | M0039 | 6.07E-06 |
| **98** | *ITM2B* | *LY86* | 0 | M0039 | 5.77E-06 |
| **99** | *ITM2B* | *LTC4S* | 0 | M0039 | 5.54E-06 |
| **100** | *ITM2B* | *YBX1* | 0 | M0039 | 5.54E-06 |
| **101** | *ITM2B* | *TREM2* | 0 | M0039 | 5.44E-06 |
| **102** | *ITM2B* | *TYROBP* | 0 | M0039 | 5.28E-06 |
| **103** | *ITM2B* | *PSME2* | 0 | M0039 | 5.23E-06 |
| **104** | *ITM2B* | *MPC2* | 0 | M0039 | 5.22E-06 |
| **105** | *ITM2B* | *PNISR* | 0 | M0039 | 5.21E-06 |
| **106** | *ITM2B* | *CDK2AP2* | 0 | M0039 | 5.13E-06 |
| **107** | *ITM2B* | *CXCL12* | 0 | M0039 | 5.00E-06 |
| **108** | *ITM2B* | *ACTB* | 0 | M0039 | 4.99E-06 |
| **109** | *ITM2B* | *NPC2* | 0 | M0039 | 4.85E-06 |
| **110** | *ITM2B* | *BIN1* | 0 | M0039 | 4.70E-06 |
| **111** | *ITM2B* | *RSRP1* | 0 | M0039 | 4.70E-06 |
| **112** | *ITM2B* | *TBXAS1* | 0 | M0039 | 4.59E-06 |
| **113** | *ITM2B* | *C1QA* | 0 | M0039 | 4.58E-06 |
| **114** | *ITM2B* | *TMBIM6* | 0 | M0039 | 4.58E-06 |
| **115** | *ITM2B* | *HYOU1* | 0 | M0039 | 4.30E-06 |
| **116** | *ITM2B* | *LGALS1* | 0 | M0039 | 4.30E-06 |
| **117** | *ITM2B* | *ZFP36L2* | 0 | M0039 | 3.96E-06 |
| **118** | *ITM2B* | *LAMP1* | 0 | M0039 | 3.95E-06 |
| **119** | *ITM2B* | *SEC61B* | 0 | M0039 | 3.60E-06 |
| **120** | *ITM2B* | *C1QB* | 0 | M0039 | 3.52E-06 |
| **121** | *ITM2B* | *RPL8* | 0 | M0039 | 3.40E-06 |
| **122** | *ITM2B* | *PRELID1* | 0 | M0039 | 3.38E-06 |
| **123** | *ITM2B* | *CYCS* | 0 | M0039 | 3.34E-06 |
| **124** | *ITM2B* | *RPLP0* | 0 | M0039 | 3.25E-06 |
| **125** | *ITM2B* | *OLFML3* | 0 | M0039 | 3.14E-06 |
| **126** | *ITM2B* | *N4BP2L2* | 0 | M0039 | 3.12E-06 |
| **127** | *ITM2B* | *HSP90AB1* | 0 | M0039 | 3.07E-06 |
| **128** | *ITM2B* | *IL1B* | 0 | M0039 | 3.05E-06 |
| **130** | *ITM2B* | *SRSF5* | 0 | M0039 | 2.93E-06 |
| **131** | *ITM2B* | *C1QC* | 0 | M0039 | 2.90E-06 |

Supplementary Table 2

| | |
|---|---|
| **1** | *FTL* |
| **2** | *TMSB4X* |
| **3** | *FTH1* |
| **4** | *ACTB* |
| **5** | *SPP1* |
| **6** | *PSAP* |
| **7** | *TPT1* |
| **8** | *CTSD* |
| **9** | *B2M* |
| **10** | *ACTG1* |
| **11** | *CTSB* |
| **12** | *CTSZ* |
| **13** | *RPS27* |
| **14** | *COL3A1* |
| **15** | *RPL37A* |
| **16** | *GPNMB* |
| **17** | *COL1A1* |
| **18** | *LGALS1* |
| **19** | *LGMN* |
| **20** | *TMSB10* |
| **21** | *CD68* |
| **22** | *RPS29* |
| **23** | *GPX1* |
| **24** | *RPL23* |
| **25** | *S100A11* |
| **26** | *SEPP1* |
| **27** | *RPLP1* |
| **28** | *RPL13AP5* |
| **29** | *C1QC* |
| **30** | *RPL38* |
| **31** | *RPS6* |
| **32** | *CD63* |
| **33** | *RPL31* |
| **34** | *RPS12* |
| **35** | *MYL6* |
| **36** | *RPS4X* |
| **37** | *RPLP2* |
| **38** | *PFN1* |
| **39** | *CFL1* |
| **40** | *FCER1G* |
| **41** | *RPL27* |
| **42** | *RPL30* |
| **43** | *NPC2* |
| **44** | *RPL19* |
| **45** | *RPS11* |
| **46** | *TYROBP* |
| **47** | *ATP5E* |
| **48** | *RPS18* |
| **49** | *COL1A2* |
| **50** | *SPARC* |
| **51** | *VIM* |
| **52** | *RPS28* |
| **53** | *RPL35* |
| **54** | *CSTB* |
| **55** | *RPS24* |
| **56** | *RPS16* |
| **57** | *LAPTM5* |
| **58** | *RPL11* |
| **59** | *C1QB* |
| **60** | *EEF1A1* |
| **61** | *IFI30* |
| **62** | *GAPDH* |
| **63** | *DLK1* |
| **64** | *RPS15A* |
| **65** | *GRN* |
| **66** | *TUBA1B* |
| **67** | *RPL35A* |
| **68** | *H19* |
| **69** | *RPS8* |
| **70** | *CD14* |
| **71** | *LIPA* |
| **72** | *RPS20* |
| **73** | *ANXA2* |
| **74** | *CD81* |
| **75** | *RPL26* |
| **76** | *ITM2B* |
| **77** | *CST3* |
| **78** | *RPS2* |
| **79** | *SOD2* |
| **80** | *FUCA1* |
| **81** | *RPL7A* |
| **82** | *GNB2L1* |
| **83** | *SAT1* |
| **84** | *OAZ1* |
| **85** | *RPL3* |
| **86** | *ATP6AP2* |
| **87** | *EEF2* |
| **88** | *RPLP0* |
| **89** | *RPS14* |
| **90** | *F13A1* |
| **91** | *ARHGDIB* |
| **92** | *CAPG* |
| **93** | *ATP6V0C* |
| **94** | *GNAS* |
| **95** | *FAU* |
| **96** | *RPL8* |
| **97** | *ASAH1* |
| **98** | *C1QA* |
| **99** | *CTSH* |
| **100** | *SDCBP* |
